# Adequate coating and change in morphology increase the performances of silver nanoparticular biocides

**DOI:** 10.64898/2026.05.11.724204

**Authors:** Bartłomiej Pawłowski, Hanna Błażyca, Jaana Huotari, Véronique Collin-Faure, Elisabeth Chartier-Garcia, Satu Salo, Elisabeth Darrouzet, Olgierd Jeremiasz, Thierry Rabilloud

## Abstract

Silver has been used as a biocide for centuries, mostly in health-oriented applications. However, as a biocide, silver is toxic not only to its intended targets, mainly bacteria and fungi, but also to all living cells. Because of this toxicity, it is desirable to use forms of silver that maximize the required biocidal activity while minimizing the amount of silver that will be released in the environment at the end of life of the product. Silver nano objects are a good compromise for such requirements. The high surface to volume ratio allows for good reactivity and thus good biocidal activity, while the small amount of silver present in nano objects allows for a limited environmental release at the product end of life. In this work, we tested three types of silver nano objects. The first type, polyvinylpyrrolidone-coated silver nanoparticles (nAg-PVP) were used as a control nanoparticle, as this type of nanoparticle is now widespread. We also manufactured and tested maltodextrin-coated silver nanoparticles (nAg-MD) and micrometric (20 µm in two dimensions and a few nanometers in the third one) silver flakes (µAgSF). For these three silver nano objects, we investigated the biocidal activity by stringent tests using both Staphylococcus aureus and Escherichia coli as target bacteria. In addition, we investigated toxicity on mammalian macrophages or keratinocytes cell lines, as well as on an insect hemocyte cell line. Our results showed that the two innovative silver nano objects (nAg-MD and even more µAgSF), showed both a better bactericidal activity and a lesser toxicity than the reference nAg-PVP nanoparticles. In addition, we also checked that beyond toxicity, the silver nano objects did not induce an inflammatory reaction, making them safer to use.

## 1. Introduction

Silver is among the oldest biocides used on a large scale by mankind, as mentioned in [1], in a medicinal use owing to its price, while cheaper copper was also used for agriculture [2]. Silver was very widely used in the pre-antibiotic era, both as an oral drug and as a topical drug, a usage that it is still common nowadays [3]. However, the extensive use of silver as a biocide in medicine came with serious drawbacks. Although usually classified as a precious metal, silver is also a non-essential, post transition metal. As such, it is poorly eliminated from living organisms and silver ion interferes with biological processes, e.g. by complexing to cysteines. This can lead to direct enzyme inhibition (e.g. in [4,5]), or to replacement of true essential metal ions in biologically important structures, such as zinc fingers [6]. These perturbations of important biological processes lead to a disease called argyria, which has been described in detail for a century or so [7–9]. Combined with the use of antibiotics, which offer a much better benefit/risk ratio, these adverse effects have confined the medicinal uses of silver to topical products, although a too intensive use of silver containing products can still lead to adverse effects [10].

Among the recent use of silver nanoparticles in topical health products, wound dressings are quite popular [11–14], although their clinical efficacy seems moderate [15,16]. One of the advantages of silver nanoparticles as a biocide, e.g. in the application of wound dressings, is the very wide scope of action of silver, as it acts on viruses, bacteria, protozoa and fungi [17,18]. The mechanisms of action of silver have been studied in detail on many different targets (e.g. in [19–23]) and globally show an impairment in the membrane functions and the production of toxic reactive oxygen species. As these mechanisms are very generic, they explain why silver nanoparticles are also toxic to mammalian cells (e.g. in [24–31]) often at concentrations quite close to those needed for a biocidal effect [32].

The severity of this lack of selectivity of silver biocides between the targeted organisms and the untargeted ones will of course depend on the applications of the silver biocides, which are quite varied (e.g. in [33,34]), mainly because of very different exposure configurations. Nevertheless, it is always interesting to increase the selectivity of the biocides, as it widens their scope of applications. For silver biocides, the particle size has been shown to be a major driver of toxicity [24,35–37], together with the shape [38,39]. In addition, silver nanoparticles being quite hydrophobic, they need to be either coated with a hydrophilic agent that will prevent their aggregation in aqueous media, or to be grafted on a polymeric matrix that will prevent their reaggregation (e.g. in silver cellulose composites [14,39]). It has been shown that the surface coating, when present, modulates the toxicity and efficacy of silver nanoparticles as biocides (e.g. in [40–43]). Thus, we decided to undertake a comparative study of silver-based particular biocides different in either the coating agent or the particles size in order to see if we could increase the selectivity between toxicity to bacteria and toxicity to mammalian cells. To this purpose, we used the same integrated strategy (both efficacy and toxicity tests conducted in parallel) than we recently used for copper-based biocides [44].

## 2. Material and methods

### 2.1. Nanoparticles synthesis

#### 2.1.1 Reagents

Analytical grade silver nitrate (AgNO3), potassium hydroxide (KOH), technical grade polyvinylpyrrolidone (MW: 30 000) were obtained from P.P.H. “STANLAB” Sp. z o.o.. Technical grade citric acid (C6H8O7 x H2O) was obtained from Biomus Sp. z o.o.. Technical grade heptahydrate of iron(II) sulfate (FeSO4 × 7H2O) was obtained from WARCHEM Sp. z o.o. Technical grade maltodextrin (C Dry MD 01910, DE = 15, DP = 8) was obtained from CARGILL. Analytical grade isopropyl alcohol, initially obtained from PureLand was continuously distilled and used within Helioenergia Facilities (overall purity = 95%). For all processes, self-produced demineralized water (conductivity < 25µS/cm) was used.

#### 2.1.2 PVP-coated silver nanoparticles (nAg-PVP)

Silver particles coated with polyvinylpyrrolidone were prepared as follows: 25.32 g of potassium hydroxide and 12.49 g of PVP were fully dissolved in 3 liters of demineralized water and stirred mechanically for 30 minutes. Meanwhile, 23.48 g of silver nitrate were dissolved in 100 milliliters of demineralized water using a glass stir rod. The silver nitrate solution was then added to the reactor containing the potassium hydroxide and PVP mixture. The entire solution was mechanically stirred for 14 hours.

Following this, mechanical stirring was halted and the solution started to sediment in approx. 60 minutes. The resulting material was separated by decantation and washed twice with acetone (100ml each time) followed by further decantation. The obtained pulp was then transferred to ceramic trays and dried at 55°C for at least 24h. The particles were then dispersed by resuspension at 10% (w/v) in distilled water followed by sonication in a water bath (ultrasound bath ULSONIX 3 × 60W) for 30 minutes. The dispersion was re-homogeneized just before use by water bath sonication for 15 minutes.

#### 2.1.2 Maltodextrin-coated silver nanoparticles (nAg-MD)

Silver particles coated with maltodextrin were prepared as follows: 21 g of potassium hydroxide and 3 g of maltodextrin were fully dissolved in 5 liters of demineralized water and stirred mechanically for 30 minutes. Meanwhile, 18 g of silver nitrate was dissolved in 2 liters of demineralized water using a glass stir rod. The silver nitrate solution was then added to the reactor containing the potassium hydroxide and maltodextrin mixture. The entire solution was mechanically stirred for 14 hours.

Following this, 7 liters of isopropyl alcohol were added to destabilize the solution, which was then stirred for 30 minutes and left to sediment. The resulting material was separated by decantation and the obtained pulp was transferred to ceramic trays. For obtaining dry silver nanoparticles, the pulp was dried at 55°C for at least 24h.

Alternatively, the obtained pulp was transferred into 3l beaker. If needed, solution was diluted with demineralized water to 1 liter. Next, the whole mixture was destabilized 5 more times (including sedimentation and decantation) with 1 liters of isopropyl alcohol per liter of initial suspension. Finally, the final water suspension was evaporated to obtain 250ml of silver dispersion. This dispersion was used directly in subsequent experiments.

#### 2.1.3. Silver microflakes (µAgSF)

Silver microflakes were prepared as follows: 880g of iron(II) sulfate heptahydrate were fully dissolved in 30 liters of demineralized water and stirred mechanically for 30 minutes. Meanwhile, 200g of silver nitrate and 30g of citric acid were fully dissolved in 10 liters of demineralized water using a glass stir rod. The acidic silver nitrate solution was then added to the reactor containing iron(II) sulfate heptahydrate solution as a slow and steady stream under mechanical stirring. Following the addition, the entire solution was mechanically stirred for 2 hours.

The mechanical stirring was then halted and the solution started to sediment in approx. 60minutes. The resulting material was separated by decantation and washed twice with demineralized water (10 liters each time) and once with isopropyl alcohol (5 liters) followed by further decantation. The obtained pulp was transferred to ceramic trays and dried at 55°C for at least 24h. The particles were then dispersed by resuspension at 10% (w/v) in distilled water followed by sonication in a water bath (ultrasound bath ULSONIX 3 × 60W) for 30 minutes. The dispersion was re-homogeneized just before use by water bath sonication for 15 minutes.

### 2.2. Nanoparticles characterization

The silver nano-objects were characterized by X-ray fluorescence (XRF) for their chemical composition, scanning electron microscopy (SEM), for their morphology and dry size and dynamic light scattering (DLS), for their in-solution behavior.

For XRF, dry silver powder particles were characterized with a Fischer X-Ray XDML instrument. A high-resolution scanning electron microscope - LEO Gemini 1530. was used for SEM

For DLS, the particle size distribution of silver particles in water dispersions was obtained with a Zetasizer Nano-S analyzer.

### 2.3. Antimicrobial activity

The Minimal Inhibitory Concentration (MIC) method was used, as described previously [44]. However, we prefer to detail the methods her again for the consistency of the publication. To this purpose, the nanopowders suspensions coming from the various syntheses were serially diluted (two-fold dilutions) in sterile milliQ water. The bacterial inoculum, grown overnight in Muller Hinton Broth at 37°C with shaking, was diluted to a concentration of approximately 5 × 10 ^6^CFU/ml. Each dilution of the nano powder was mixed with an equal volume of bacterial suspension and incubated at 37°C for 20 hours. The experiments were performed in microwell plates, with each well containing 0.2 ml and three replicates per sample. To prevent drying out during incubation, the plates were placed in a plastic container with water at the bottom. Following the incubation period, microbial survival was determined by culturing 10 µl of each dilution on Plate Count agar and incubating at 37°C for 24 hours. The test bacteria used were *Staphylococcus aureus* VTT E-70045 and *Escherichia coli* VTT E-94564 (VTT Culture Collection, VTT Technical Research Centre of Finland Ltd, Espoo, Finland).

### 2.4. Nanotoxicology

In this section, we used the same methods than those used previously on copper-based biocides [44]. However, we provide here a complete methods section for the readers convenience.

#### 2.4.1. Cell survival assay

To evaluate the toxicity of the nanoparticles, cell survival assays were performed, using the MTT reduction method [45]. As target cell lines, we used the J774A.1 murine macrophage cell line [46] and the HaCat human keratinocyte cell line [47]. The J774A.1 cell line was purchased from ECACC (Salisbury, UK), while the HaCat cell line was a gift of Pr N. Fusenig (Heidelberg, Germany). Both cell lines were cultures in DMEM supplemented with 10% fetal bovine serum.

For routine culture, the J77A.1 cells were seeded at 200,000 cells/ml in non adherent cell culture flasks (Greiner) and harvested two day later at 800,000-1,000,000 cells/ml. The HaCat cells were seeded in adherent culture flasks at 20,000 cells/cm2 and recovered at confluence 3 days letter. For passaging the cells, the cell layer was rinsed three times in serum-free DMEM, and then incubated in trypsin EDTA (0.25% trypsin, 1mM EDTA in PBS) at a ratio of 0.05ml/cm2 at 37°C for 5-10 minutes. The reaction was stopped by diluting the cell suspension in the trypsin solution into 5 volumes of complete culture medium (with serum). The cell suspension was then centrifuged (200g, 5 minutes) to pellet the cells, which were then resuspended in complete culture medium, numerated using the Trypan blue method [48] and diluted for culture.

For the survival assay tests, the cells were seeded in 24 wells adherent culture plates at a density of 500,000 cells/ml for J774A.1 cells and 80,000 cells/cm2 for HaCat cells. After 24 hours, the nanoparticles suspensions were added to the wells and the cells returned to the cell culture incubator for 24 hours. At the end of the exposure period, the culture medium was removed, the cells layers were rinsed once with DMEM without phenol red, and then incubated for 1 hour at 37°C in serum-free and phenol red-free DMEM containing 0.2mg/ml MTT (predissolved at 20mg/ml in ethanol). At the end of the MTT exposure period the culture medium was removed and the produced formazan eluted in a solution containing 90% ethanol and 1% acetic acid in water (all by volumes). The plates were placed on a plate shaker for 30 minutes at room temperature. The eluate was transferred to a fresh plate to avoid interferences due to the opacity of the added nanoparticles, and the absorbance at 570 nm was measured by a BMG Fluostar Omega plate reader.

In order to better mimic occupational exposure, a repeated exposure scheme was also used. In this scheme, the cells were seeded in culture medium containing 1% horse serum instead of 10% fetal bovine serum to limit cell proliferation [49]. After 24 hours, a split dose, i.e. a quarter of the final cumulated dose was added to the medium. After 24 hours of exposure the medium was changed and a split dose was added. After 24 hours another split dose was added. Finally the medium was changed and a fourth split dose was added, before a final 24 hours culture period. The medium was then removed and the cell viability testing protocol was carried out.

#### 2.4.3. Cytokine release assays

In order to evaluate the pro-inflammatory secretion of cytokines induced by the nanoparticles, the cell supernatants collected after the acute exposure to the nanoparticles were collected, centrifuged at 15,000g for 30 minutes to remove the nanoparticles, and then assayed for the production of Interleukin 6 (IL-6) and Tumor Necrosis Factor alpha (TNF), was measured. The chemokines MCP1 (for J774A.1 cells) and Interleukin 8 (IL-8), for HaCat cells were also measured. The cytokine bead array method was used for the multiplexed measurements of the cytokines, using the Human Inflammatory Cytokine kit (BD biosciences, #551811 for HaCat cells and the Cytometric Bead Array Mouse Inflammation Kit (catalog numbers 558299, 558301 and 558266, BD Biosciences) for J774A.1 cells.

#### 2.4.4. Phagocytosis assay

This test was performed only on J774A.1 cells, as they are known to be phagocytic. After acute exposure to the nanoparticles, the cells were exposed for 3 hours at 37°C to latex beads (carboxypolystyrene, yellow-green labelled, 0.5µm, #15700 from Polysciences). The cells were rinsed twice with PBS, and analyzed for the green fluorescence (excitation 488nm, emission 527/32 nm) on a Melody flow cytometer. Controls with no latex beads showed no green fluorescence.

#### 2.4.5. Toxicity on insect cells

The toxicity test was performed on Schneider S2 cells, which are a model for insect hemocytes [50]. For culture maintenance, the cells (purchased from DSMZ, Germany) were grown at 25°C in T75 flasks in an air atmosphere and in a culture medium composed of 4.5 parts of Schneider medium, 4.5 parts of TC100 medium and 1 part of fetal bovine serum. Cells were routinely seeded at 40,000/80,000 cells/cm2 and harvested by scraping after 2 to 4 days.

For the toxicity testing, the cells were seeded at 250,000 cells/cm2 in 24 well culture plates. After 24 hours of settling and growth, the cells were treated with varying concentrations of the biocides for 24 hours. After this incubation period, the cell viability was assessed with a WST1 viability kit (# 11644807001 from Sigma Aldrich Millipore, Saint Quentin Fallavier, France) with a dilution factor of 1:10 and an incubation period of 4 hours. At the end of the incubation period, the plates were centrifuged (200g for 5 minutes) and the supernatant collected for analysis by absorbance at 440 nm. The cell layer was then fixed with 50% ethanol for 40 minutes, and the ethanol was removed. The cell layer was then rehydrated in culture medium overnight, and a second WST1 test was performed to determine the assay background, which was then subtracted from the cell assay to obtained the corrected absorbance.

## 3. Results

### 3.1. Nanoparticles characterization

We first characterized the purity of the silver nano-objects by X-ray diffraction. The results, shown in Figure 1, suggest that silver is the only metal present. We did not detect any other metallic element, such as adsorbed iron in the microflakes preparation.

**Figure 1.**
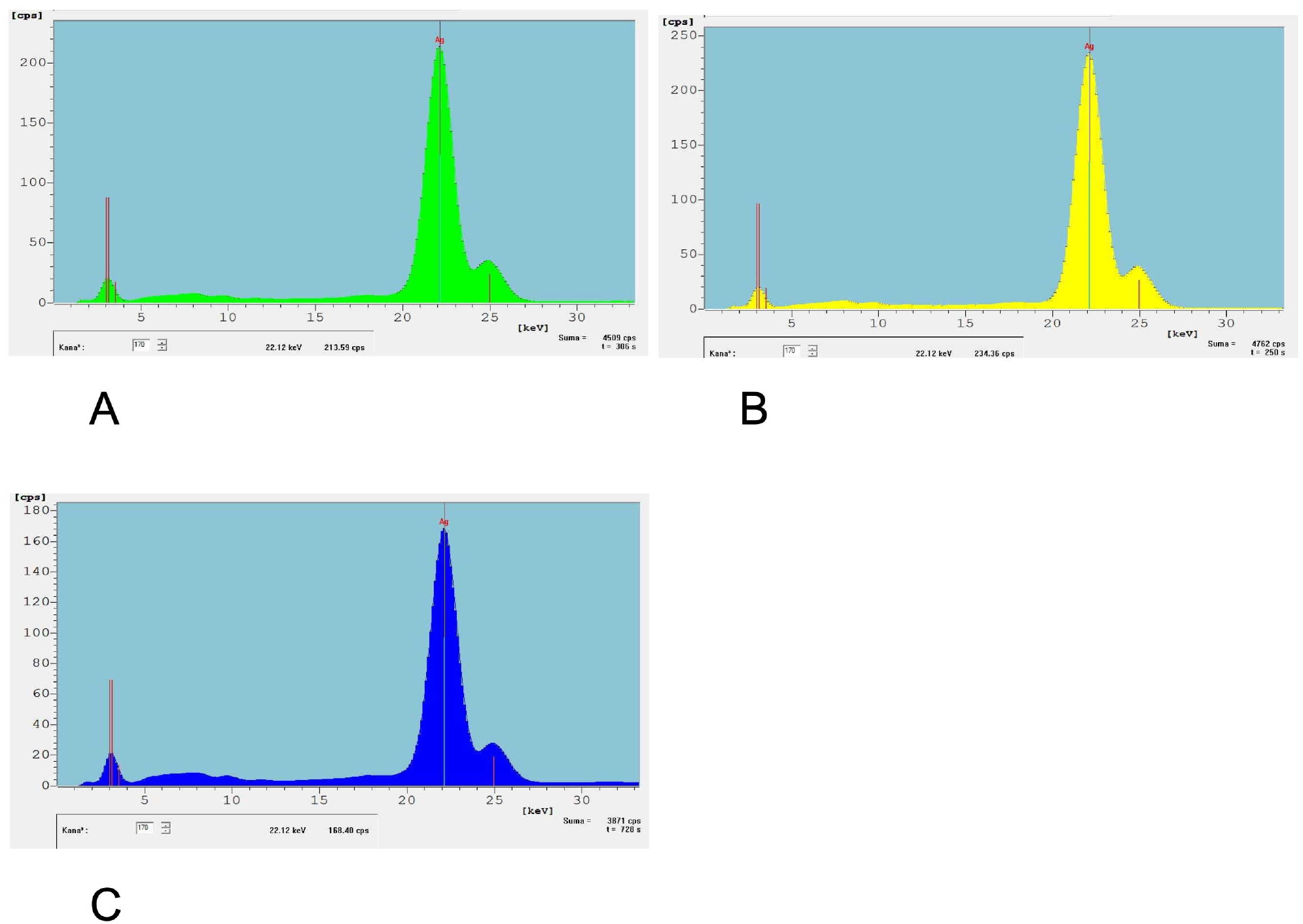
X-ray diffraction diagrams of the three nano-objects A: diagram for nAg-PVP B: diagram for nAg-MD C: diagram for µAgSF

We then characterized the morphology of the silver nano-objects by SEM. In the images shown in Figure 2, nAg-PVP and nAg-MD appeared in SEM as fused primary nanoparticles. These primary particles were in the 30-40nm range for the nAg-PVP nanoparticles, and in the 20-40nm range for the nAg-MD nanoparticles. Silver microflakes showed a completely different morphology, and appeared as flakes that were only a few nanometers thick, but micrometric in their other two dimensions, with a modal size of 10-20µm, with the occurrence of smaller fragments.

**Figure 2.**
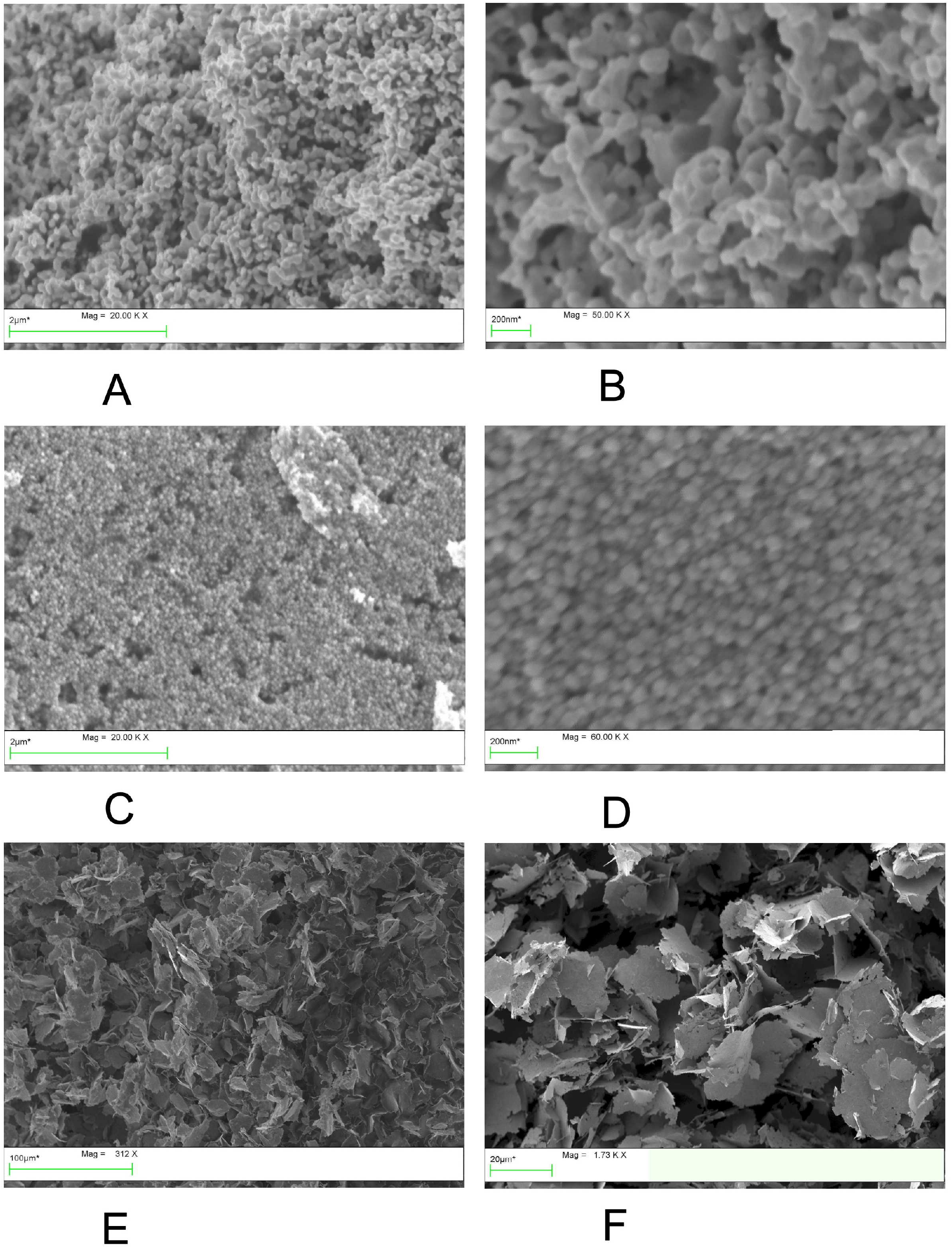
SEM images of the three nano-objects. The magnifications are indicated for each image A, B: images for nAg-PVP C, D: images for nAg-MD E, F: images for µAgSF

Finally, as it is a key parameter for the biological experiments, we characterized the mean in solution characteristics of the silver nano-objects dispersions by DLS. The results, displayed in Figure 3, showed that nAg-PVP and nAg-MD consisted of two populations. In the case of nAg-PVP, the minor (10% in intensity) population showed a mean hydrodynamic diameter of 14±4 nm, while the major population (90% in intensity) showed a mean hydrodynamic diameter of 116±75 nm. In the case of nAg-MD, the minor (14% in intensity) population showed a mean hydrodynamic diameter of 12±3.5 nm, while the major population (90% in intensity) showed a mean hydrodynamic diameter of 68±33 nm. Finally, µAgSF showed a single population with a mean hydrodynamic diameter of 161±36 nm. This low value may thus represent only the smallest population, which does not sediment quickly.

**Figure 3.**
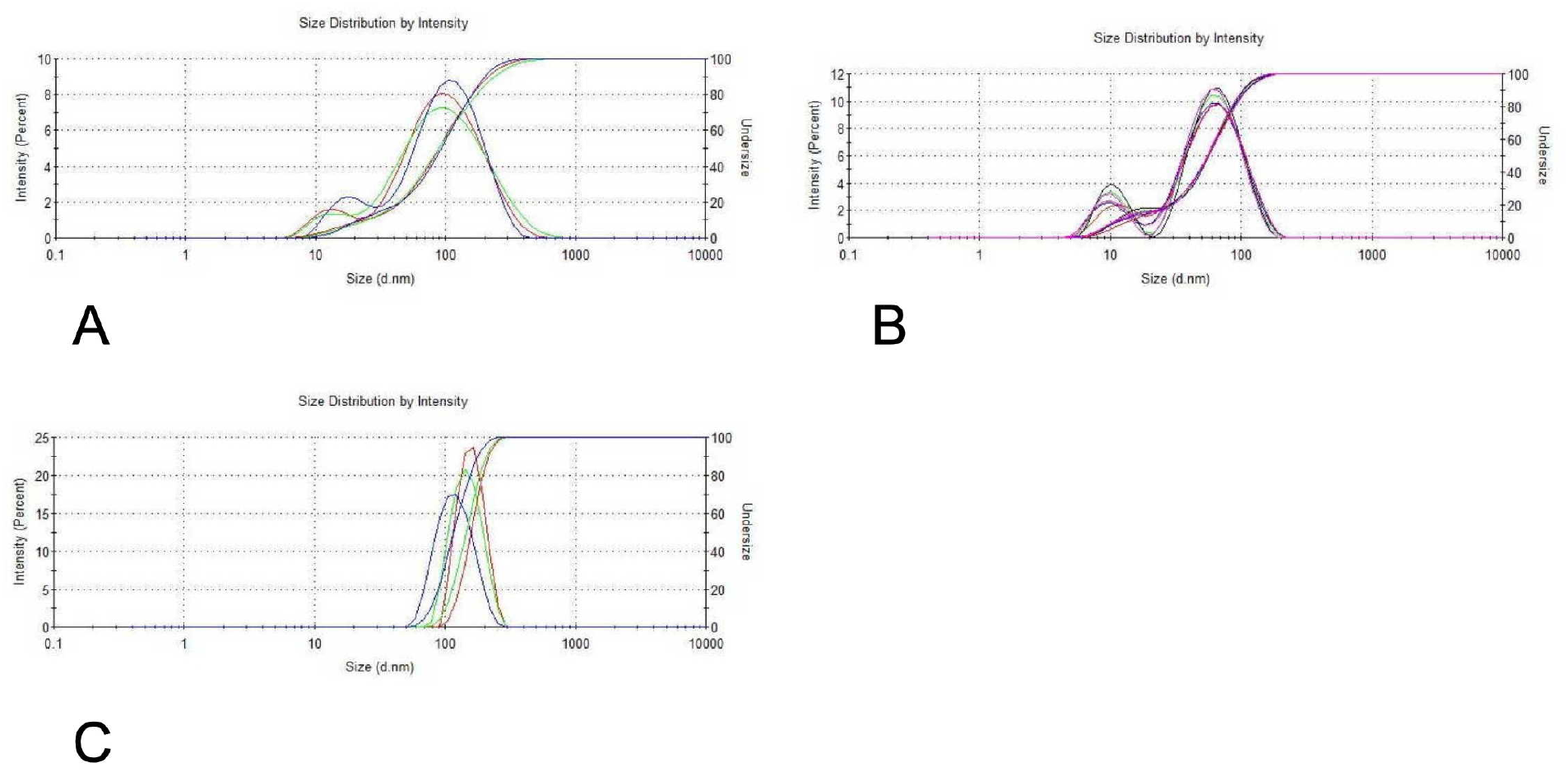
DLS diagrams of the three nano-objects A: diagram for nAg-PVP B: diagram for nAg-MD C: diagram for µAgSF

### 3.2. Antibacterial efficacy

The serial dilution MIC method was used to determine the biocides antibacterial efficacy (Table 1). Three independent experiments were performed, and their individual results are presented

**Table 1:**
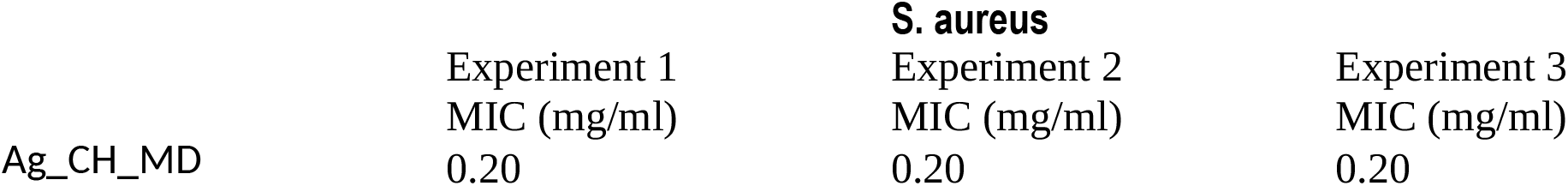
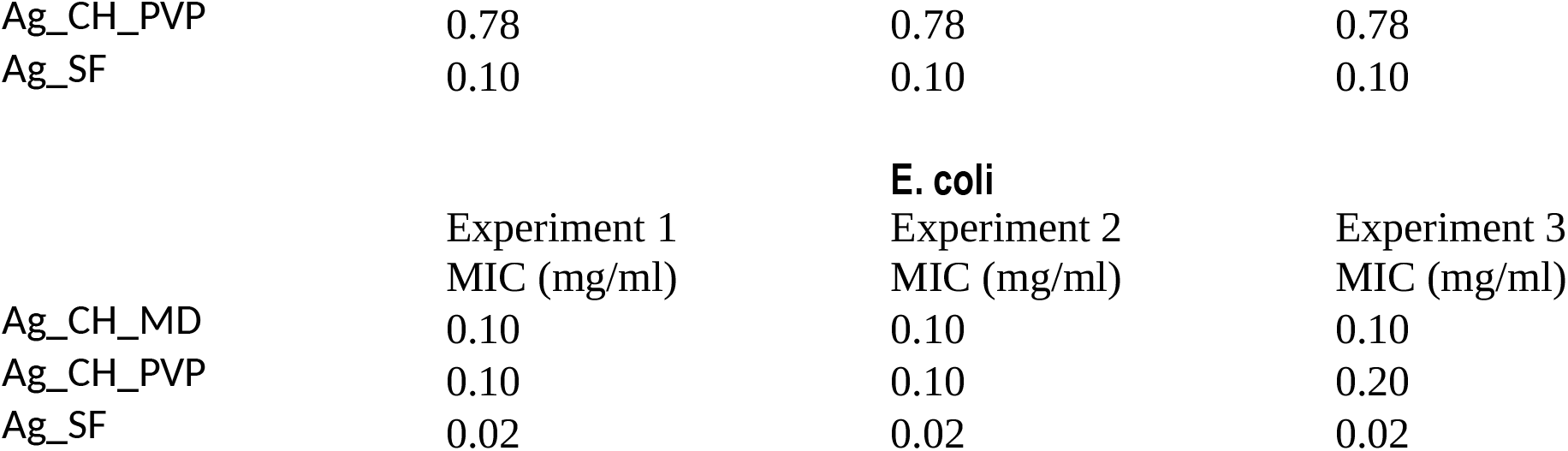
MIC for the silver-based nanoparticles.

### 3.3. Biological effects on mammalian cells

We used two different cell lines to investigate the potential toxic effects of the biocides on mammalian cells. As macrophages are the primary scavenger cells taking care of all kinds of particulate materials in our bodies, whatever the contamination route, we used the J774A.1 mouse macrophage cell line. Furthermore, as cutaneous exposure is a major expected contamination route, we also used the HaCat human keratinocyte cell line.

We first investigated the effects on cell viability in an acute exposure mode (24 hours). The results, displayed on Figure 4, showed first a very low toxicity for the microsilver flakes, in both cell systems. Regarding the classical nAg-PVP particles, they showed a greater toxicity for macrophages (LD50= 7.5 µg/ml) than for keratinocytes (LD50=75µg/ml), which can be explained by the phagocytosis of the nanoparticles agglomerates by the macrophages, keratinocytes being non-phagocytic. Maltodextrin coated silver nanoparticles (nAg-MD) showed an intermediate toxicity. The LD50 was not reached at 200µg/ml in these initial experiments on macrophages, while a lower LD50 (180µg/ml) was found for keratinocytes.

**Figure 4.**
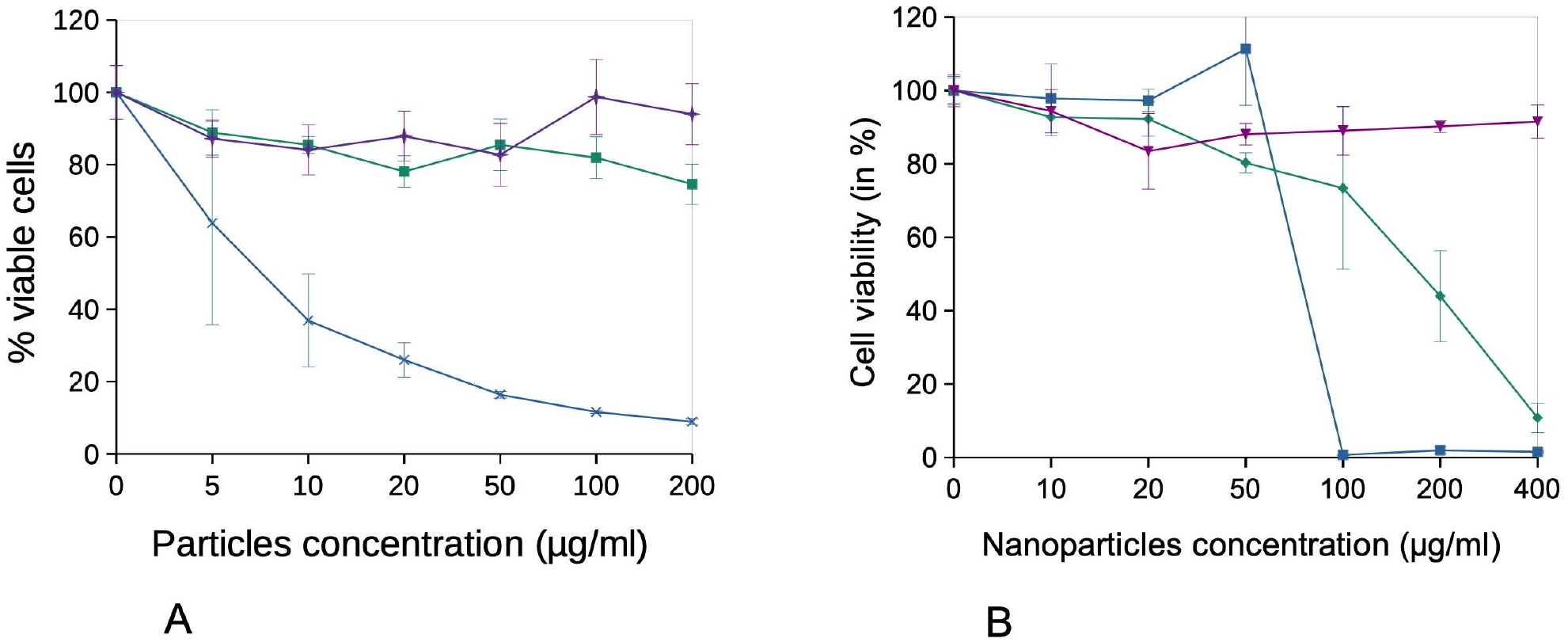
Survival curves of cells treated for 24 hours with silver nanoparticles J774A.1 cells (panel A) and HaCaT cells (panel B) were treated at confluence for 24 hours with silver nanoparticles in complete cell culture medium (containing 10% FBS). The cell viability was then measured by the MTT assay. The results are displayed as mean ± standard deviation (N=4). Blue curve: cells exposed to nAg-PVP Green curve: cells exposed to nAg-MD Purple curve: cells exposed to µAgSF

We then investigated the toxic effects of the silver biocides in a repeated exposure mode, where the total dose is fractionated in four daily doses. This scheme was selected to better evaluate the effects of the silver particles on workers, which may be exposed repeatedly to sub-toxic doses during their weekly working days. The results, displayed on Figure 5, showed contrasted results. For nAg-PVP, the toxicity was similar in both exposure modes for macrophages. For keratinocytes, the toxicity in the acute exposure mode was higher in this serum-low medium than in the serum rich medium used in the initial experiments. Moreover, the toxicity was somewhat inferior in the repeated exposure scheme. For the nAg-MD particles, the higher toxicity under acute exposure and serum-low medium was found again for both cell types. However, for this particle, the repeated exposure mode led to a marked decrease in toxicity, especially for keratinocytes. A lesser toxicity in the repeated mode can be attributed to the fact that cells exposed to sub-toxic doses are able to build a protective response against further exposure.

**Figure 5.**
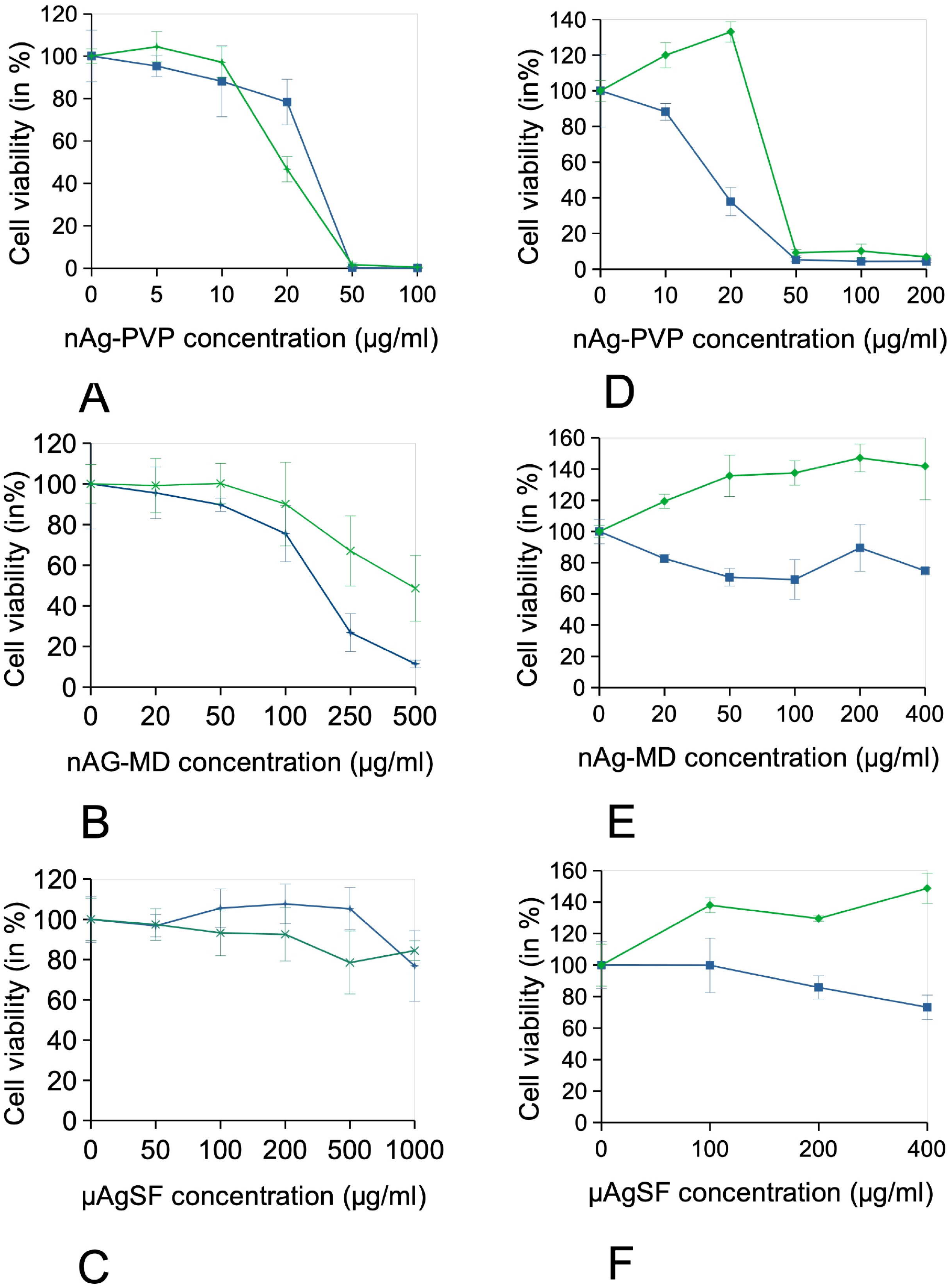
Survival curves of cells treated for 24 hours with silver-based nanoparticles Survival curves of cells treated for 4 days with copper oxide-based nanoparticles J774A.1 cells (panels A-C) and HaCaT cells (panels D-F) were treated at confluence either for 24 hours or 4 days with copper oxide-based nanoparticles in adapted cell culture medium (containing 1% horse serum). The indicated dose is the cumulated exposure concentration. The cell viability was then measured by the MTT assay. The results are displayed as mean ± standard deviation (N=4). Blue curve: acute exposure Green curve: repeated exposure

Apparent viabilities above 100% were measured, as for µAgSF. This just translates the fact that particles-exposed cells perceive them as a stress and increase their metabolism (the parameter measured by the MTT test) to cope with this stress. If the particle do not induce cell mortality, this phenomenon appears as an apparent over-viability. Finally, µAgSF appeared as very weakly toxic.

The examples of silicosis and asbestosis show that particles can induce adverse effects by inflammation-driven mechanisms [51]. We thus assayed the secretion of pro-inflammatory cytokines and chimiokines by cells treated with non-toxic concentrations of the particles. The results, displayed on Tables 2 and 3, showed no induction of the secretion of interleukin 6 and sometimes even a significant decrease, opposite to what has been described on the J774A.1 line for amorphous silica, for example [52].

**Table 2:**
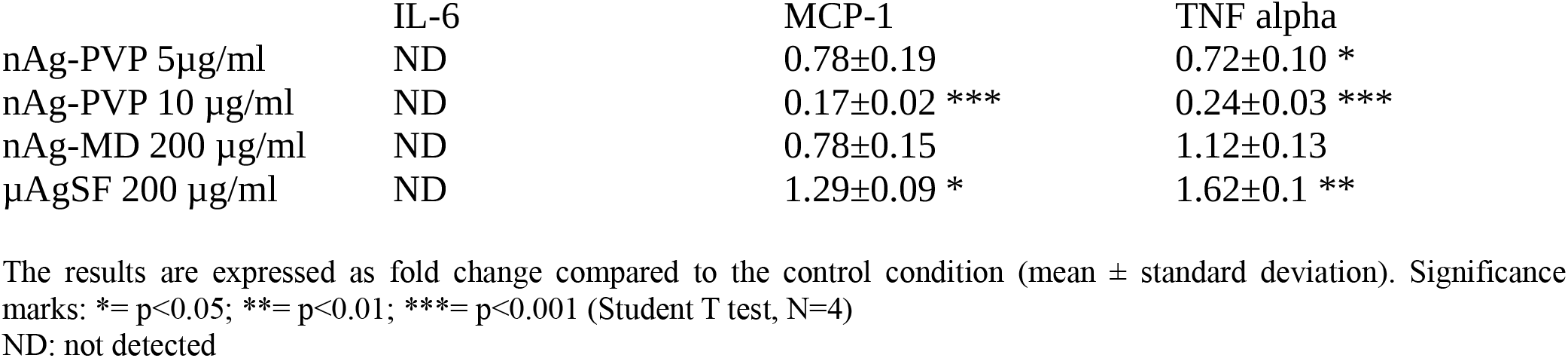
cytokine secretion in J774A.1 cells exposed to nanoparticles.

**Table 3:**
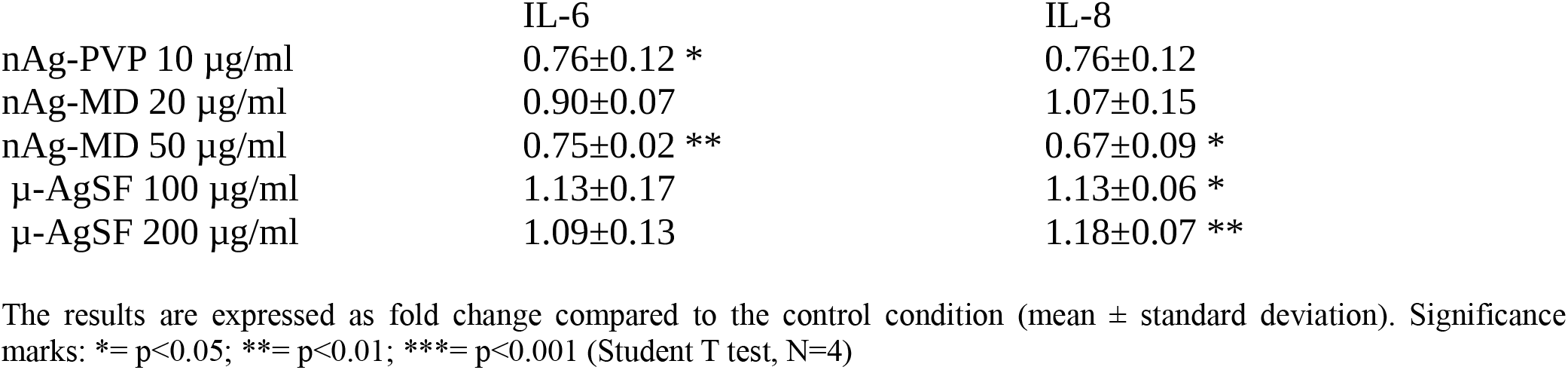
cytokine secretion in HaCaT cells exposed to nanoparticles.

A decrease, sometimes significant, was also observed for the MCP-1 chemokine except for µAgSF for which a moderate but significant increase was observed. The situation was different for TNF, where a small (ca. 1.2-fold) but significant increase was observed in response to nAg-MD and µAgSF particles, while a decrease was observed in response to nAg-PVP. However, this increase is much lower compared to the one observed in response to amorphous silica [52].

For HaCaT cells, we measured the human chemokine interleukin 8 in lieu of MCP-1, and IL-6. TNF secretion could not be detected in any of the tested conditions.

No significant increase in the secretion of IL-6 was observed. Regarding IL-8, a moderate but significant increase was observed for µAgSF.

Phagocytosis being an important macrophage function, e.g. in the clearance of pathogens, we investigated the potential impact of non toxic concentrations of the silver biocides on this biological parameter. The results, displayed on Figure 6, showed a slight (one third) but significant decrease in the phagocytic capacity of J774A.1 cells in response to µAgSF, while nAg-MD induced a more important decrease (-50 to -75%).

**Figure 6.**
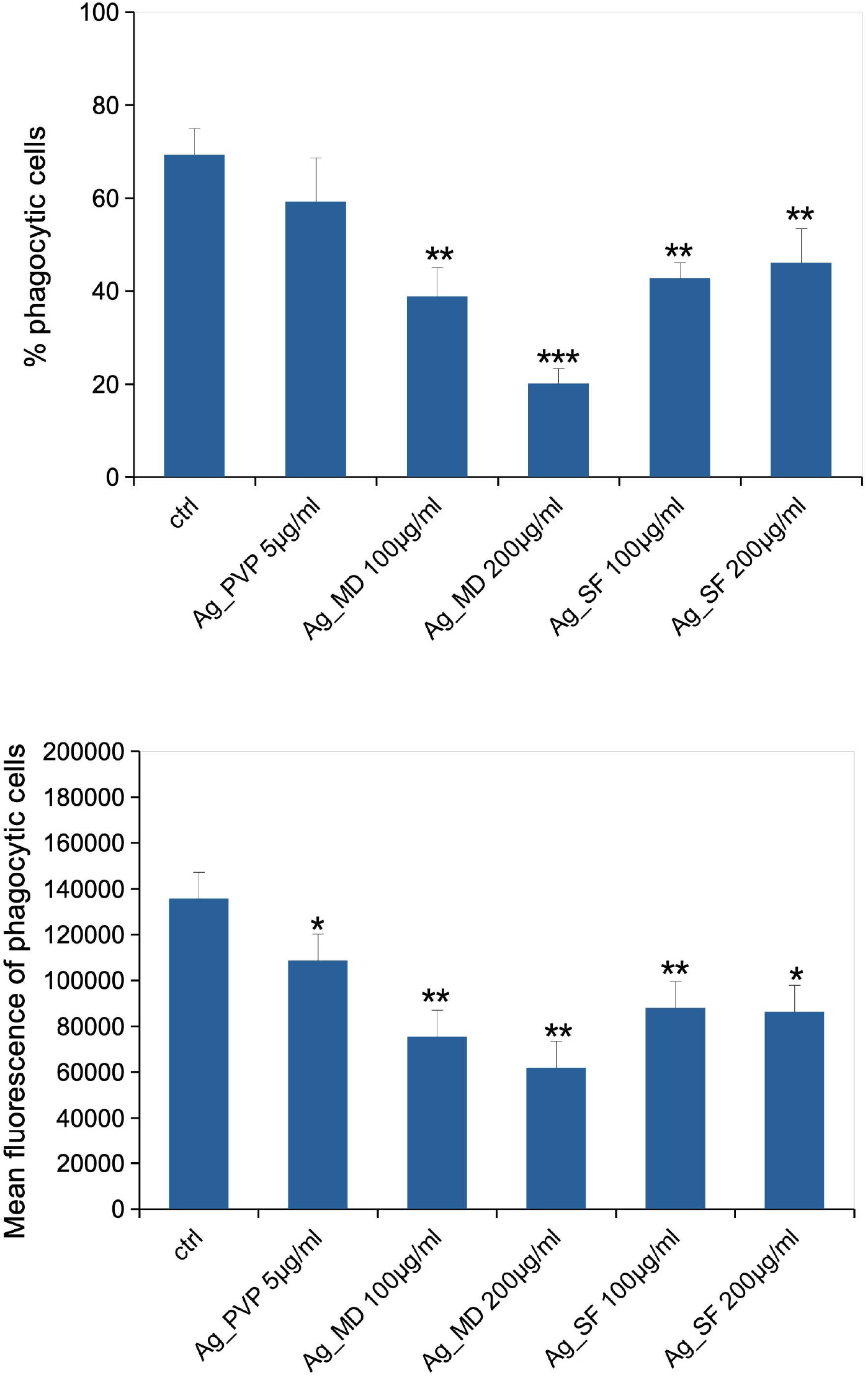
Phagocytic activity of J774A.1 cells treated for 24 hours with silver nanoparticles In the top panel, the proportion of phagocytosis positive cells (i.e. cells having internalized at least one latex bead in three hours) is shown. The bottom panel shows the amount of fluorescence due to beads internalization within the three hours phagocytosis window. Results are shown as mean ± standard deviation. Significance mark: *=p<0.05 (Student T test, N=4)

### 3.4. Biological effects on insect cells

Finally, we tested the effect of the silver biocides on Schneider cells, which are a model of insect hemocytes, i.e. the equivalent of vertebrates macrophages [50]. The results, displayed on Figure 7 showed an interesting bias of the metabolic test. Indeed, an apparent increase in viability is observed at low, non toxic concentrations. As we performed a background test on killed cells having internalized the particles, we can exclude a direct redox activity of the silver particles. By definition, viability cannot exceed 100%. As we worked at cellular confluence in the flasks, this result can only be explained by the fact that internalization of the silver particles requires energy consumption by the cells, thereby increasing the metabolism and thus the signal in the metabolic test used as a viability test. Of note, this effect was also detected, although to a lesser extent, in HaCat cells (see above). Nevertheless, we could detect a toxic response at higher concentrations for nAg-PVP and nAg-MD, with respective LD50 of 80 µg/ml and 600 µg/ml. No LD50 could be determined for µAgSF at practical concentrations.

**Figure 7.**
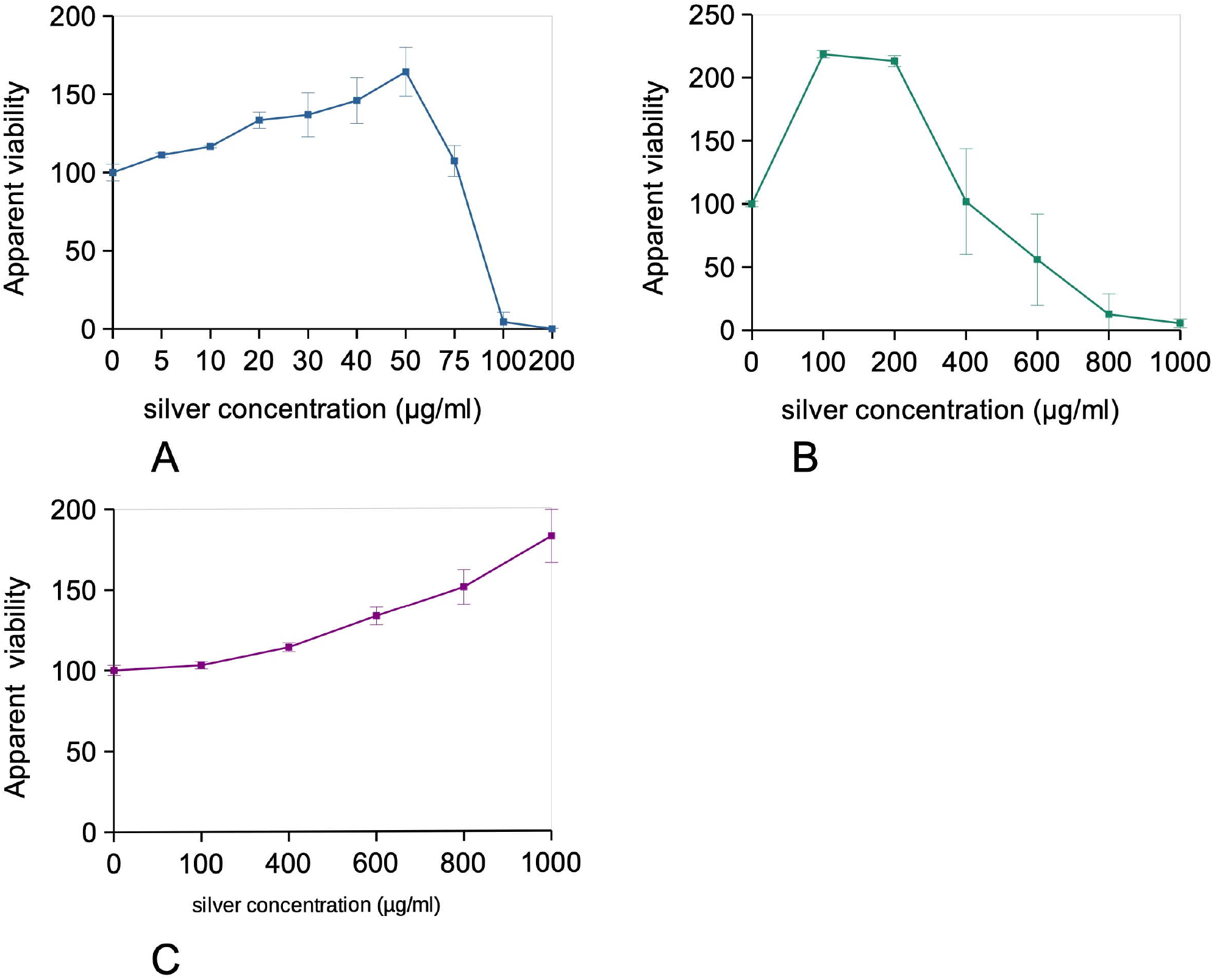
Viability tests of cells treated for 24 hours with silver particles Schneider cells were treated at confluence for 24 hours with silver particles in complete cell culture medium. The cell viability was then measured by the WST1 assay. The results are displayed as mean ± standard deviation (N=4). A: cells treated with nAg-PVP B: cells treated with nAg-MD C: cells treated with µAgSF

## Discussion

As for every biocide, the main challenge is to maximize the contrast between the biocidal effect on targeted organisms (i.e. efficacy) and the biocidal effects on untargeted organisms (i.e. toxicity). However, this differential parameter has been seldom explored for silver-based biocides. There is on the one side an important corpus of scientific literature exploring the antibacterial efficacy of silver biocides (reviewed for example in [53]) leaving out of scope the toxicity issues, and on the other side an important corpus of scientific literature focused on the toxic effects (reviewed for example in [54]) but leaving out of scope the antibacterial efficacy parameter. However, there are a few papers exploring at the same time both areas for the very same silver nanoparticles (e.g. in [36,55–60]). Within them, most are devoted to so-called green synthesized silver nanoparticles, i.e. nanoparticles produced with completely uncharacterized plant extracts [55–59], and only a few dealt, as we do in the present paper, with silver nanoparticles produced with controlled chemicals [36,41,60].

A further complication arises from the fact that different publications use different methods to characterize the antibacterial efficiency. For example in the publication dealing silver-PVP nanoparticles [60], i.e. a particle type present in this paper too, the authors use a disk diffusion method, used in a few other papers (e.g. in [61]), and not the serial dilution method, which we used in our tests and which is the most widespread one (e.g. in [41,42,55,57]). Nevertheless, we can try to compare the data. Hamed et al. found a MIC of 0.1 mg/ml against S. aureus and 0.2 mg/ml against E. coli [60], to be compared to the 0.13 mg/ml against E. coli and 0.78 mg/ml against S. aureus found in our study. Although direct comparison is further complicated by the fact that different bacterial strains are used in different publications, these concentrations fall in the same order of magnitude. Regarding the cytotoxicity of their silver-PVP nanoparticles, Hamed et al. found a LD50 close to 200µg/ml on hepatocytes [60], while we found a LD50 close to 75µg/ml for keratinocytes and <10µg/ml for macrophages, i.e. a phagocytic cell type that will internalize much more particles than non phagocytic types such as fibroblasts, keratinocytes or hepatocytes. Thus, our data agree with the data of Hamed et al. to characterize silver-PVP as a biocide with relatively low contrast between efficacy and toxicity. Citrate-coated silver NPs, which dissolve more rapidly than polymer-coated silver NPs, proved more toxic to both bacteria and mammalian cells [36].

A way to alter positively the contrast between toxicity and efficacy would be to change the coating agent covering the silver nanoparticles. Indeed, this solution has been explored in the literature and has been found to be efficient on at least one of the two parameters, especially when the capping ligand either contains sulfur, which has a high affinity for silver [42], or when the capping ligand is a biological polymer (e.g. in [41,43]). This path is also followed in silver nanoparticles synthesized with complex biological extracts (e.g. in [55–59]). However, the natural variability of such complex extracts may result in variable results over time in different batches, making this avenue more complex to implement for industrial processes.

We thus followed the path of changing the ligand for another, chemically controlled one, and synthesized maltodextrin-coated silver nanoparticles. Compared to dextran used in previous work [41,43], maltodextrin is a much shorter polymer. The grade that we used has a polymerization degree (DP) of 8, meaning that the average chain length is 8 glucose unit, to be compared to the >200 glucose units of the dextran 40 used by Ferreira et al. [43], and the > 1000 glucose units of the dextran 250 used by Yang et al. [41].

Although this ligand was relatively simple, its effects on the performances of the biocides were important. One the one hand the MIC decreased and reached 0.2 mg/ml for S. aureus and 0.1 mg/ml for E. coli. More importantly, the LD50 for mammalian cells increased considerably and was higher than 100µg/ml, i.e. in the range of the antibacterial concentrations. Dextran-coated silver nanoparticles also showed increased biocidal performances compared to PVP-coated silver nanoparticles [41]. However, they were still quite toxic to cells, with a LD50 at 24 hours of exposure close to 10µg/ml [41].

Another avenue that we explored to increase the efficacy/toxicity contrast was to play with the shape and size of the particles. The antibacterial effects of the silver nanoparticles are linked to both extracellular dissolution and silver ion diffusion, as exemplified by experiments using the disk diffusion method [61] and to direct particle-cell contact [62]. This explains why small nanoparticles, which show both a higher penetration in complex matrices and a higher surface/volume ratio, leading to increased dissolution at constant silver mass, induce stronger biocidal effects [36]. The toxic mechanisms toward mammalian cells are somewhat different, as they depend mainly on intracellular dissolution after particles internalization [63]. This explains why small silver nanoparticles are more toxic to mammalian cells than bigger ones [36], but it also opens an avenue to decrease toxicity toward mammalian cells. Indeed, even phagocytic cells show an upper limit to the size of the particles that they can internalize, which is close to 10-15 µm [64]. Besides of size, shape also plays a role, and at equal size, discoidal objects are less internalized than spherical ones [65]. We thus wondered if we could use these properties and manufactured silver flakes of micrometric size. Opposite to spherical silver microparticles which would afford a poor surface/volume ratio and thus poor efficacy coupled with high silver consumption for a given effect, flakes afford a high surface/volume ratio and thus may be able to couple high dissolution and or contact with bacterial cells, coupled with low internalization in mammalian cells and thus low toxicity. In fact this avenue has been indirectly explored in clay-silver nanoparticles composites [66], where the low size of the nanoparticles brings efficient dissolution while the large size of the clay flakes brings low internalization.

Our results showed that this avenue was fruitful. The observed MIC reached 100 µg/ml for S. aureus and 20µg/ml for E. coli, while the LD20 for mammalian cells was not reached at 200 µg/ml and appeared at 400 µg/ml or even above. Compared to the silver-clay nanocomposite system [66], the silver flakes are made in one step and there is no risk of particles detachment upon use, which may release free silver NPs. We also verified that the silver flakes did not induce inflammatory reactions, such as those induced by long silver nanowires (e.g. in [67]). In fact the only interference that we detected was a slight decrease in phagocytosis, which was observed for all three silver nanomaterials tested.

In conclusion, our results show that beside the classical avenue of altering the coating to change the contrast between efficacy and toxicity, playing with the aspect ratio seems to be an even more promising avenue to increase this efficacy/toxicity. However, we need to be careful when playing with this parameter. One-dimensional high aspect ratio objects, such as fibers and wires, are often very toxic by chronic inflammation mechanisms, as shown by the infamous examples of asbestosis and silicosis [68,69]. These mechanisms are based on a peculiar mechanism called frustrated phagocytosis [70]. However, this mechanism does not work with two-dimensional high aspect ratio objects such as plates, as exemplified by our current results on silver microplates and by previous knowledge on shape influence on particles internalization by animal cells [65]. Thus, it appears that playing with the shape parameter along these lines could be an interesting avenue of research not only for silver-based biocides, as exemplified here, but also for biocides based on other metals or metal oxides.

